# Synergistic effects of anionic surfactants on coronavirus (SARS-CoV-2) virucidal efficiency of sanitizing fluids to fight COVID-19

**DOI:** 10.1101/2020.05.29.124107

**Authors:** Reza Jahromi, Vahid Mogharab, Hossein Jahromi, Arezoo Avazpour

## Abstract

Our surrounding environment, especially often-touched contaminated surfaces, plays an important role in the transmission of pathogens in society. The shortage of effective sanitizing fluids, however, became a global challenge quickly after the coronavirus disease-19 (COVID-19) outbreak in December 2019. In this study, we present the effect of surfactants on coronavirus (SARS-CoV-2) virucidal efficiency in sanitizing fluids. Sodium dodecylbenzenesulfonate (SDBS), sodium laureth sulfate (SLS), and two commercial dish soap and liquid hand soap were studied with the goal of evaporation rate reduction in sanitizing liquids to maximize surface contact time. Twelve fluids with different recipes composed of ethanol, isopropanol, SDBS, SLS, glycerin, and water of standardized hardness (WSH) were tested for their evaporation time and virucidal efficiency. Evaporation time increased by 17-63% when surfactant agents were added to the liquid. In addition, surfactant incorporation enhanced the virucidal efficiency between 15-27% according to the 4-field test in the EN 16615:2015 European Standard method. Most importantly, however, we found that surfactant addition provides a synergistic effect with alcohols to inactivate the SARS-CoV-2 virus. This study provides a simple, yet effective solution to improve the virucidal efficiency of commonly used sanitizers.

## 1. Introduction

The coronavirus pandemic, also known as the COVID-19 pandemic, is an ongoing pandemic of coronavirus disease 2019, caused by severe acute respiratory syndrome coronavirus 2 (SARS-CoV-2) [1]. To date, more than 4.5 million cases have been confirmed, with more than 300,000 deaths globally. Surfaces can become contaminated by hands, objects, settling of virus-containing aerosols, or contaminated fluids [2, 3]. Therefore, these surfaces may play an important role in the transmission of pathogens in society [4, 5]. Manual disinfection of surfaces by wiping and spraying antibacterial fluids is an important part of the healthcare setting. The shortage of effective sanitizing liquids, however, became a global challenge quickly after the COVID-19 outbreak in December 2019. Ethanol and isopropanol are the major active ingredient (~70-80%) of most common sanitizers, but the high volatility of these alcohols makes them less effective when sprayed on surfaces. The alcohol retention time on the surface plays a key role in its viricidal efficiency. A common solution to overcome this challenge is to include less volatile material (such as aloe vera gel) in sanitizer formulation. However, while blended with gels, the final solution becomes thicker and more difficult to spray on surfaces.

Anionic surfactants that are used in the manufacturing of liquid soaps and detergents, are also known for their antibacterial efficiency because of their dual functionality. The hydrophobic side of these surfactants can dissolve the outer layer of viruses and bacteria, while the hydrophilic side dissolves in water [6, 7]. Therefore, these agents act as an emulsifier to remove the contamination. After the COVID-19 pandemic, world health organization (WHO) has recommended that individuals wash hands more frequently and at least for 20 seconds to ensure bacterial and virus removal from the skin [8, 9]. It has been shown that the lipid outer layer of SARS-CoV-2 disrupts after long enough contact time with anionic surfactants [10].

In this study, we explored the effect of anionic surfactants addition to sanitizing liquids with different recipes. The main objective of this study was to reduce the volatility of the sanitizers without negatively influencing their thickness (viscosity) while boosting their virucidal properties. Two well known anionic surfactants, sodium dodecylbenzenesulfonate (SDBS) and sodium laureth sulfate (SLS) that are commonly used in dish soaps and liquid soaps, respectively, were studied at 3% concentration. In addition, commercial dish soap and liquid hand soap were subjected to experimentation with the goal of providing a simple, yet effective solution for daily applications. All tested liquids were also studied for their SARS-CoC-2 viricidal efficiencies.

## 2. Experiments

The SARS-CoV-2 coronavirus was obtained from Molecular Epidemiology Laboratory at Shiraz University of Medical Science, Iran. Ethanol, isopropanol, glycerin, SDBS, and SLS were obtained from Millipore Sigma (Beijing, China). Two commercial dish soap and hand soap were purchased from Golrang (Tehran, Iran). Water of standardized hardness (WSH) was used in control experiments and also as a diluting agent in the formulation of sanitizing fluids. Twelve sanitizing fluids with different recipes, as shown in Table 1, were prepared to examine the effect of individual components and mixtures on evaporation rate and SARS-CoV-2 virucidal efficiency of the solutions.

**Table 1:** Sanitizing solution recipes, labels, and evaporation times.

| Composition<br>(wt. %) | Solution label |  |  |  |  |  |  |  |  |  |  |  |  |
| --- | --- | --- | --- | --- | --- | --- | --- | --- | --- | --- | --- | --- | --- |
|  | WSH | S1 | S2 | S3 | S4 | S5 | S6 | S7 | S8 | S9 | S10 | S11 | S12 |
| SDBS | 0 | 0 | 0 | 0 | 0 | 3 | 3 | 0 | 0 | 0 | 0 | 0 | 0 |
| SLS | 0 | 0 | 0 | 0 | 0 | 0 | 0 | 3 | 0 | 0 | 0 | 0 | 0 |
| Ethanol | 0 | 70 | 0 | 35 | 35 | 70 | 70 | 0 | 0 | 70 | 35 | 0 | 0 |
| Isopropanol | 0 | 0 | 70 | 35 | 35 | 0 | 0 | 70 | 70 | 0 | 35 | 0 | 0 |
| Hand soap | 0 | 0 | 0 | 0 | 0 | 0 | 0 | 0 | 3 | 0 | 0 | 0 | 3 |
| Dish soap | 0 | 0 | 0 | 0 | 0 | 0 | 0 | 0 | 0 | 3 | 3 | 3 | 0 |
| Glycerin | 0 | 0 | 0 | 0 | 3 | 0 | 3 | 0 | 0 | 0 | 3 | 0 | 0 |
| WSH | 100 | 30 | 30 | 30 | 27 | 30 | 27 | 27 | 27 | 27 | 24 | 97 | 97 |

For evaporation tests, PVC material with polyurethane (PUR) surface coating was sliced into 20×50 cm (2 mm thickness) pieces and placed on a flat bench, as shown in Figure 1. Sanitizing fluids were sprayed three times (from a distance of ~30 cm from the surface) on the PVC object. Fluid evaporation was recorded using a Sony HXR-NX100 camera, and videos were processed using Ulead Videostudio software. A 60-watt amber light was installed 100 cm above the surface to shine light on the surface to record better resolution videos. The surrounding environment was isolated to eliminate possible air convection, and the room temperature was maintained at ~27 °C. The exact amount of evaporation time was recorded for each sprayed liquid.

**Figure 1:**
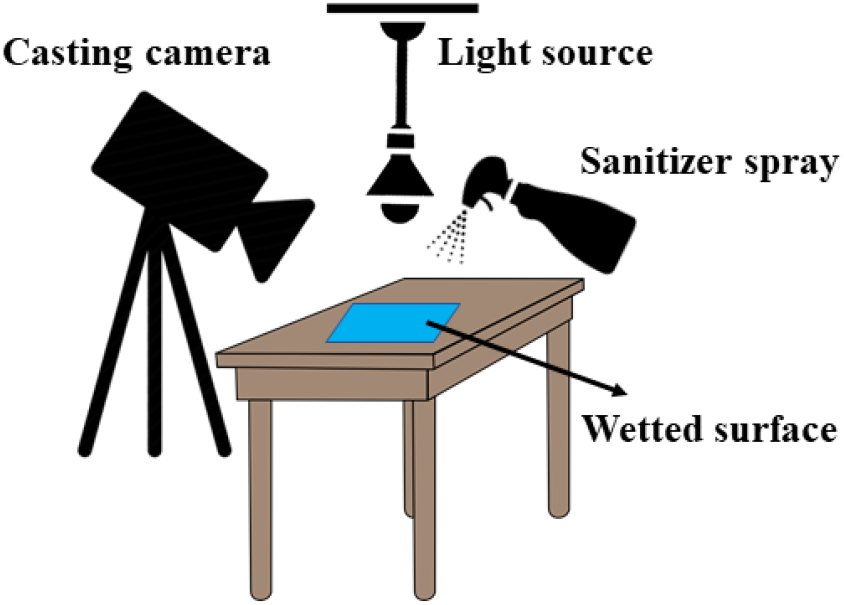
Schematics of the evaporation test experiments.

The coronavirus suspension was prepared by infecting monolayers of A549 cell (human lung epithelial carcinoma cells) lines. The virus titers of these suspensions ranged from 10^5^ to 10^10^ TCID_50_/ml. To determine the virucidal efficiency of each liquid, the 4-field test was performed according to EN 16615:2015 European Standard with slight modification [11]. Briefly, four squares as test fields were marked on the PVC with PUR surface coating material figuring a row, 5 cm away from each other. The marked test field 1 on this flooring was inoculated with the SARS-CoV-2 virus suspension. 10 μl inoculum was pipetted on the first test field and distributed with a spatula. A granite block was rapidly moved from test field 1 to test field 4 and back within no longer than 5 seconds. After 10 min contact time and drying, the sanitizing fluid was sprayed on the contaminated surface three times and allowed for 1 min. The coronavirus was then recovered from all four fields with a nylon swab. Swabs of each field were transferred to 5 ml Minimum Essential Medium (MEM), respectively, and tubes were vortexed for 120 s. Virus titres were determined by endpoint dilution techniques according to the EN 14476:2015 European Standard and calculated using the method of Kärber and Spearman [12, 13]. The virus reduction factor (RF) on the test fields was then determined according to the method described in details by Becker et al. [14]. All experiments were carried out in triplicates.

## 3. Results and discussion

Water-based sanitizing solutions normally consist of ~70% alcohol (ethanol and/or isopropanol) as an active ingredient, ~3% glycerin to reduce dehydration if applied to the skin, and a small amount of fragrance. WSH, tested as control liquid, showed 97 s evaporation time using the method described above. To study the evaporation of sanitizing fluids, individual and multiple-compound tests were performed. Evaporation test results are presented in Figure 2. The evaporation time of 70% isopropanol solution (S2) was higher than that of ethanol solution (S1) that was ascribed to a higher molecular weight of isopropanol than ethanol. The effect of glycerin addition to a mixture of 35% ethanol and 35% isopropanol (S3 versus S4 solutions) also showed an increase in evaporation time by approximately 23% from 30 to 37 s. However, it is desired to maximize the liquid contact time with the surface to enhance the virucidal efficiency of the sanitizer. SDBS surfactant, which is one of the main components of most commercial dish soaps, significantly increased the evaporation time from 24 s (of ethanol solution S1) to 33 s (for solution S5) (~37% increase). In addition, the incorporation of 3% SLS (a major constituent of liquid hand soaps) increased the evaporation time of isopropanol solution (S2) from 35 s to 41 s in solution S7 (~17% increase). These positive results encouraged us to investigate commercial dish soap and liquid hand soaps. The volatility reduction in the case of these soaps were even more significant than those of SDBS and SLS. Notably, liquid hand soap (solution S8) increased the evaporation time of isopropanol solution by about 57% from 35 to 55s. Furthermore, the addition of 3% dish soap to the ethanol solution (S1) increased the evaporation time by about 63% from 24 to 39 s (of fluid S9), as shown in Figure 2. Solutions S11 and S12 were prepared to record the evaporation time in the absence of alcohols and glycerin. We observed that dish soap and hand soap increased the evaporation time by 9% and 15%, respectively when compared with the control experiment (WSH). Since evaporation experiments showed promise, we performed virucidal efficiency experiments thereafter.

**Figure 2:**
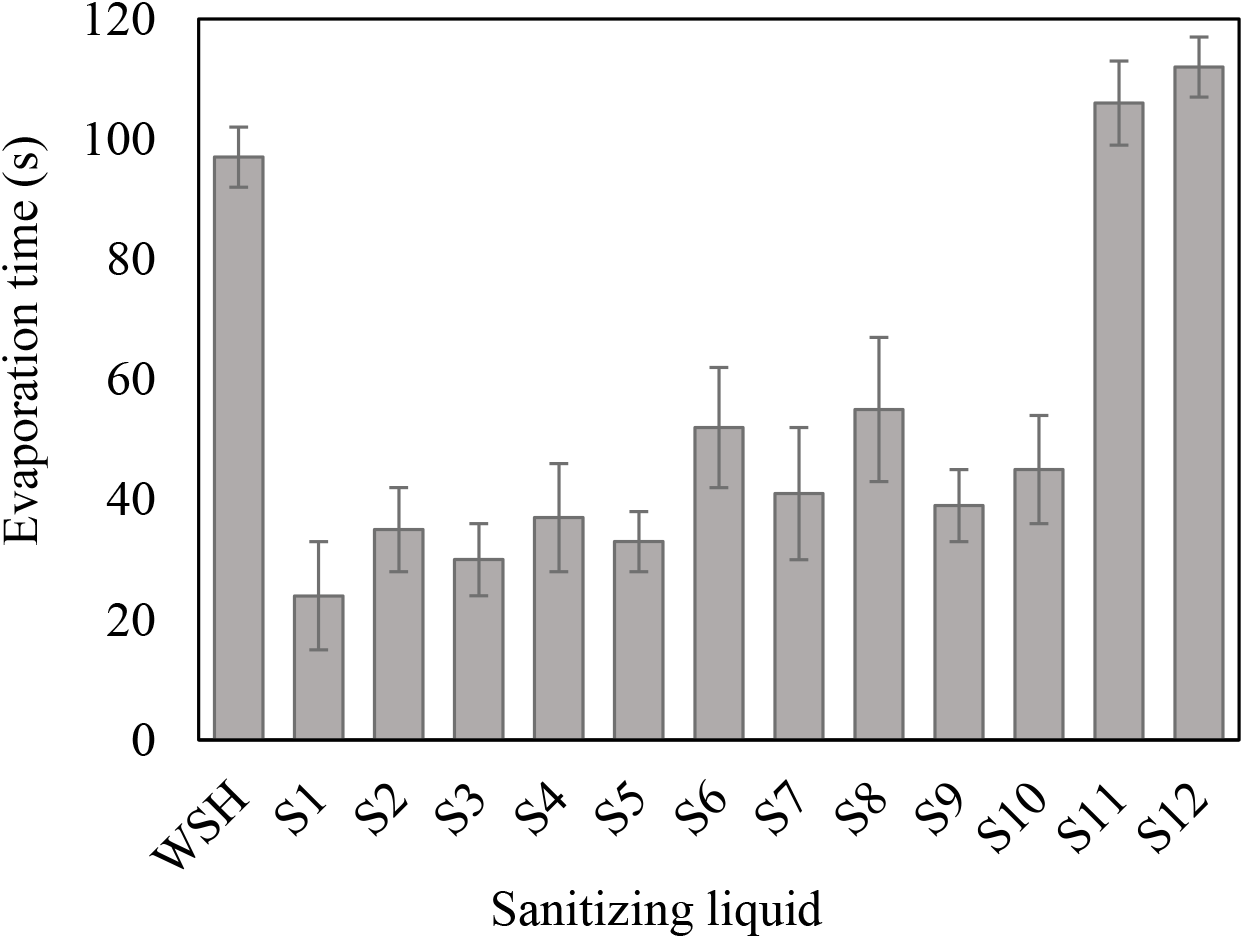
Evaporation times at 27°C (error bars show the standard deviation of three measurements)

The twelve liquids exhibited different virucidal efficacies against the chosen coronavirus (Figure 3). Isopropanol showed a slightly higher (~7%) reduction factor than ethanol solution (S1 and S2 in Figure 3). However, the combination of the alcohols (S3) did not exhibit RF between S1 and S2. The virucidal efficiency of S3 was ~13% greater than the expected value (average of S1 and S2) that could be ascribed to a different virucidal activity in S3. The addition of 3% glycerin did not influence the RF significantly as can be seen from the RF value obtained for S3 (6.2) versus S4 (6.0). This slight decrease might be a result of interference in the virucidal activity sourced from glycerin. SDBS and SLS surfactants both showed a significant positive influence on the RF of ethanol and isopropanol. As such, SDBS (in S5) increased the virucidal activity of ethanol solution by ~21%. Also, even though 3% glycerin was introduced, SDBS still increased the RF value from 6.4 of S6 to 6.6 of fluid S6. The combination effect of glycerin and SDBS is not clear at this time and requires further investigation. SLS surfactant, when added to isopropanol solution (recipe S7), exhibited ~19% increase in virucidal properties of the solution (S2 compared with S7). Fluids S8 and S9 were prepared to examine the influence of real dish soap and hand soap on RF value. Interestingly, we observed higher virucidal efficiency in the case of commercial soaps. The RF increased by ~15% when SLS was replaced with liquid hand soap (S7 versus S8). Moreover, the recorded RF for the fluid with 3% dish soap (S9) was ~16% greater than the one with 3% SDBS (S5). The excellent increase in both evaporation time and virucidal efficiency in the presence of the tested anionic surfactants and liquid soaps clearly suggested a synergistic effect between alcohols, surfactants, and virucidal activity mechanisms. Because the combination of the two alcohols (S3) demonstrated higher RF than individual alcohols (S1 and S2), only fluid S3 was tested for virucidal property in the presence of glycerin. However, the addition of dish soap to solution S4 caused a significant increase in RF by ~27% from 6 to 7.6. This result was in contrast with the former observation when the addition of 3% glycerin caused a slight decrease in fluid S4 compared with solution S3. Therefore, there could be another positive synergistic effect between the three agents (alcohol, soap, and glycerin) that creates a need for further investigation. We hypothesize that alcohols participate in the virucidal activity by dehydration mechanism, while anionic surfactants inactivate the virus via cell disruption mechanism. This hypothesis, with emphasis on the synergistic influence, will be studied as the next phase of the present investigation. Solutions S11 and S12 were prepared to explore the virucidal properties of the commercial liquid soaps, regardless of alcohols. As expected, although these components slightly increased the RF value, the changes were negligible when compared with the control experiment (WSH). Among tested fluids, recipe S8 demonstrated the greatest virucidal efficiency (RF= 7.8). This fluid consisted of 70% isopropanol, 3% hand soap, and 27% WSH. This high level of RF could, to some extent, be attributed to the presence of antibacterial agents (i.e., triclosan) in liquid hand soaps.

**Figure 3:**
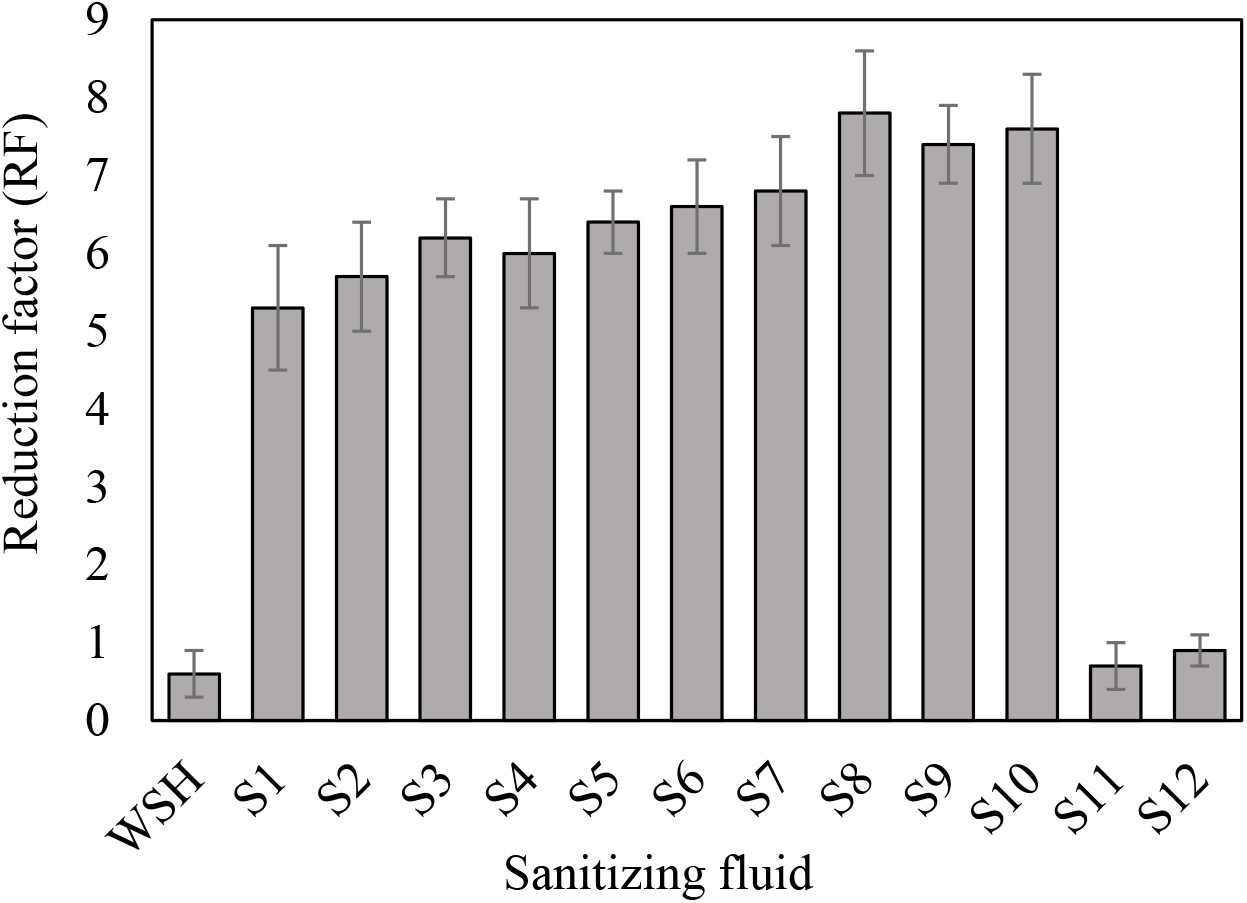
Virucidal properties of sanitizing fluids (error bars show the standard deviation of three measurements)

## 4. Conclusion

The production and use of sanitizing fluids emerged as a critical need after the COVID-19 pandemic in 2019. In this work, we explored the influence of anionic surfactants (that are the main constituents of liquid soaps) on room-temperature evaporation and virucidal efficiency of twelve fluids with different recipes. The fluids consisted of ethanol, isopropanol, dodecylbenzenesulfonate (SDBS), sodium laureth sulfate (SLS), glycerin, liquid hand soap, dish soap, and water of standardized hardness (WSH) at different ratios. The addition of surfactant agents in all experiments exhibited a significant increase in evaporation time as well as virus inactivation properties. Fluid S8 (consisted of 70% isopropanol, 3% hand soap, and 27% WSH) showed the greatest virucidal efficiency (RF= 7.8) and evaporation time (55 s). The virucidal efficiency and evaporation time of the fluids that were made of only alcohols (in WSH) or only surfactant (in WSH) were higher than the control experiment. However, the presence of all ingredients (alcohols, soap, and glycerin) increased the RF value much greater than what was anticipated. Our records clearly suggested a synergistic effect that could be attributed to different virucidal activity mechanisms. This study can provide the ongoing global challenge with a very simple solution to enhance surface disinfection efficiency to lessen the spread of SARS-CoV-2 from often-touched contaminated surfaces.

## Acknowledgment

The experimental works for this study were carried out at Jahrom University of Medical Sciences, Iran, as part of the rapid response act to the COVID-19. No human and/or animals were subjected to experimentation during this study.

